# Susceptibility of white-tailed deer (*Odocoileus virginianus*) to SARS-CoV-2

**DOI:** 10.1101/2021.01.13.426628

**Authors:** Mitchell V. Palmer, Mathias Martins, Shollie Falkenberg, Alexandra Buckley, Leonardo C. Caserta, Patrick K. Mitchell, Eric D. Cassmann, Alicia Rollins, Nancy C. Zylich, Rendall W. Renshaw, Cassandra Guarino, Bettina Wagner, Kelly Lager, Diego G. Diel

## Abstract

The origin of severe acute respiratory syndrome coronavirus 2 (SARS-CoV-2), the virus causing the global coronavirus disease 19 (COVID-19) pandemic, remains a mystery. Current evidence suggests a likely spillover into humans from an animal reservoir. Understanding the host range and identifying animal species that are susceptible to SARS-CoV-2 infection may help to elucidate the origin of the virus and the mechanisms underlying cross-species transmission to humans. Here we demonstrated that white-tailed deer (*Odocoileus virginianus*), an animal species in which the angiotensin converting enzyme 2 (ACE2) – the SARS-CoV-2 receptor – shares a high degree of similarity to humans, are highly susceptible to infection. Intranasal inoculation of deer fawns with SARS-CoV-2 resulted in established subclinical viral infection and shedding of infectious virus in nasal secretions. Notably, infected animals transmitted the virus to non-inoculated contact deer. Viral RNA was detected in multiple tissues 21 days post-inoculation (pi). All inoculated and indirect contact animals seroconverted and developed neutralizing antibodies as early as day 7 pi. The work provides important insights into the animal host range of SARS-CoV-2 and identifies white-tailed deer as a susceptible wild animal species to the virus.

**IMPORTANCE:** Given the presumed zoonotic origin of SARS-CoV-2, the human-animal-environment interface of COVID-19 pandemic is an area of great scientific and public- and animal-health interest. Identification of animal species that are susceptible to infection by SARS-CoV-2 may help to elucidate the potential origin of the virus, identify potential reservoirs or intermediate hosts, and define the mechanisms underlying cross-species transmission to humans. Additionally, it may also provide information and help to prevent potential reverse zoonosis that could lead to the establishment of a new wildlife hosts. Our data show that upon intranasal inoculation, white-tailed deer became subclinically infected and shed infectious SARS-CoV-2 in nasal secretions and feces. Importantly, indirect contact animals were infected and shed infectious virus, indicating efficient SARS-CoV-2 transmission from inoculated animals. These findings support the inclusion of wild cervid species in investigations conducted to assess potential reservoirs or sources of SARS-CoV-2 of infection.

## Introduction

Severe acute respiratory syndrome coronavirus 2 (SARS-CoV-2) is a novel coronavirus, within the genus *Betacoronavirus* (subgenus *Sarbecovirus*) of the family *Coronaviridae*, that causes coronavirus disease 2019 (COVID-19) in humans (1). COVID-19 was first reported in Wuhan, Hubei province, China in December 2019 (2). The early clusters of the disease in humans had an epidemiological link to the Huanan Seafood Wholesale market in Wuhan, where several live wild animal species were sold (2–4). Genome sequence analysis of SARS-CoV-2 revealed a high degree of similarity to coronaviruses circulating in bats (3, 5, 6), with current evidence pointing to horseshoe bats as the most likely source of the ancestral virus that crossed the species barrier to cause the global COVID-19 pandemic in humans (7, 8).

Other pathogenic zoonotic coronaviruses, including severe acute respiratory syndrome coronavirus (SARS-CoV) and Middle East respiratory syndrome coronavirus (MERS-CoV) are also believed to have originated in bat reservoirs. However, there is no evidence of direct bat-to-human transmission, and current data indicate that human infections with SARS-CoV and MERS-CoV resulted from interactions with intermediate animal hosts, such as palm civet cats (*Paguma larvata*) and dromedary camels (*Camelus dromedaries*), respectively (9–11). The epidemiological link of the first reported human cases of COVID-19 with the Huanan animal market in Wuhan, suggest that SARS-CoV-2 may have spilled over into humans from an animal host (3–5, 11–13). Early studies proposed that pangolins (*Manis* sp.) may have served as an intermediate host for SARS-CoV-2, as they are a natural reservoir for SARS-CoV-2-like coronaviruses (14). However, phylogenetic analyses and amino acid sequence analysis of the S gene of SARS-CoV-2 did not support the hypothesis of the virus arising directly from the closely related pangolin betacoronaviruses (15). A better understanding of the host range and species susceptibility of SARS-CoV-2 is critical to elucidate the origin of the virus and to identify potential animal reservoirs and routes of transmission to humans.

The tropism and host range of coronaviruses is largely determined by the Spike (S) glycoprotein, which binds to host cell receptors triggering fusion and virus entry into susceptible cells (16).

The SARS-CoV-2 S protein binds host cells through the angiotensin-converting enzyme 2 (ACE2) protein receptor (17). Comparison of the human ACE2 protein to that of over 400 vertebrate species demonstrated that the ACE2 protein of several animal species presents a high degree of amino acid conservation to the human protein (18). Further analysis of the ACE2/S binding motif and predictions of the SARS-CoV-2 S-binding propensity led to the identification of several animal species with an ACE2 protein with a high binding probability to the SARS-CoV-2 S protein (18). Not surprisingly, the majority of species with the highest S/ACE2 binding propensity are non-human primates (18). Notably, the ACE2/S protein binding motif of three species of deer, including Père David’s deer (*Elaphurus davidianus*), reindeer (*Rangifer tarandus*), and white-tailed deer (*Odocoileus virginianus*) share a high homology to the human ACE2 (18). These observations suggest a putative broad host range for SARS-CoV-2, however, the susceptibility of most of these animal species to SARS-CoV-2 infection remains unknown. Natural SARS-CoV-2 infections have been reported in dogs, cats, mink, tigers and lions in Hong Kong, Netherlands, China and the United States (19–22). The increased interest in the virus host range, in understanding the array of susceptible animal species, and the need to develop reliable animal models for SARS-CoV-2 infection, led to several experimental inoculations in domestic and wild animal species. Experimentally infected non-human primates, ferrets, minks, cats, dogs, raccoon dogs, golden Syrian hamsters, and deer mice have displayed mild to moderate clinical disease upon SARS-CoV-2 infection (23–28). Whereas experimental inoculation of swine, cattle, poultry, and fruit bats have shown that these species are either not susceptible to SARS-CoV-2 or that inoculation on these studies did not result in productive infection and sustained viral replication (23, 29–31). Here we assessed the susceptibility of white-tailed deer to SARS-CoV-2. Viral infection, clinical outcomes, shedding patterns and tissue distribution were evaluated. Additionally, transmission of SARS-CoV-2 to indirect contact animals was also investigated.

## Results

### Susceptibility of deer cells to SARS-CoV-2 infection and replication

The susceptibility of deer cells to SARS-CoV-2 infection and replication were assessed *in vitro*. Deer lung (DL) cells were inoculated with SARS-CoV-2 isolate TGR/NY/20 (32) and the ability of SARS-CoV-2 to infect these cells was compared to Vero-E6 and Vero-E6/TMPRRS2 cells. Virus infection and replication were assessed by immunofluorescence (IFA) staining using a SARS-CoV-2 N-specific monoclonal antibody. As shown in Fig. 1A, SARS-CoV-2 N expression was detected in DL cells at 24 h post-inoculation (pi), indicating that these cells are susceptible to SARS-CoV-2 infection. Importantly, staining for ACE2 confirmed expression of the SARS-CoV-2 receptor in DL cells (Fig. 1A).

**Figure 1.**
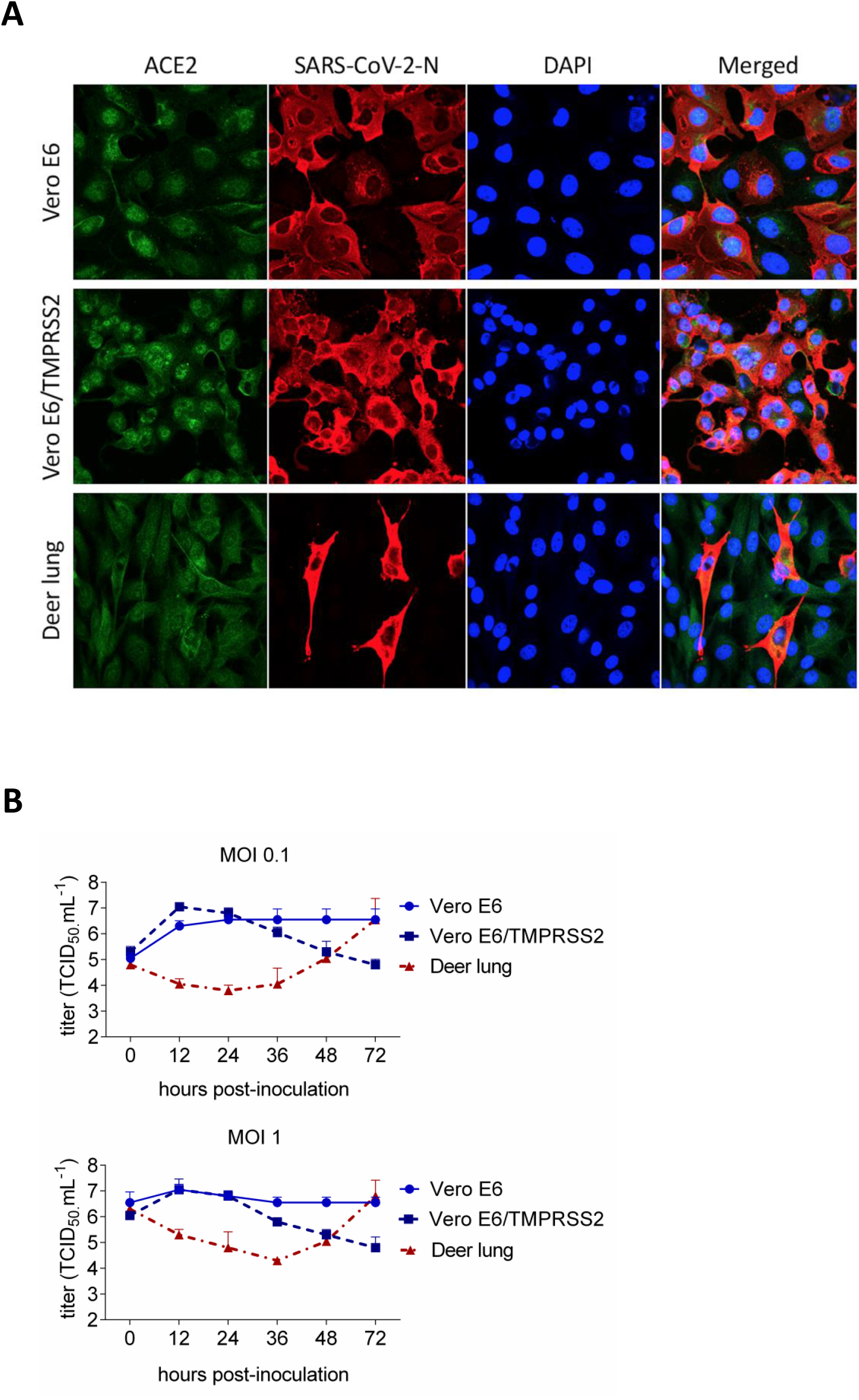
Susceptibility and replication properties of the SARS-CoV-2 in deer cells *in vitro*. (A) Deer lung (DL), Vero E6 and Vero E6/TMPRSS2 cells were inoculated with SARS-CoV-2 at a multiplicity of infection of 1 (MOI = 1). At 24 h post-inoculation (h pi), cells were fixed and subjected to an immunofluorescence assay using a monoclonal antibody (MAb) anti-ACE2 (green), and with a MAb anti-SARS-CoV-2-nucleoprotein (N) (red). Nuclear counterstain was performed with DAPI (blue). 40x magnification. (B) To assess the kinetics of replication of SARS-CoV-2, DL, Vero E6, Vero E6/TMPRSS2 were inoculated with SARS-CoV-2 isolate TGR/NY/20 (MOI = 0.1 and MOI = 1) and harvested at various time points post-inoculation (12, 24, 36, 48 and 72 h pi). Virus titers were determined on each time point using end-point dilutions and the Spearman and Karber’s method and expressed as TCID_50_.ml^-1^. Results represent the average of three independent experiments.

The replication kinetics of SARS-CoV-2 was investigated in DL cells. For comparison we included Vero-E6 and Vero-E6/TMPRSS2 cells, which are known to support efficient SARS-CoV-2 replication (33, 34). All cells were inoculated with SARS-CoV-2 at a multiplicity of infection (MOI) of 0.1 and 1, and harvested at 0, 12, 24, 36, 48 and 72 h pi. Consistent with previous studies showing efficient replication of SARS-CoV-2 in Vero-E6 and Vero-E6/TMPRSS2 cells (33, 34), SARS-CoV-2 replicated well in these cells, reaching peak titers (∼10^6^ to 10^7^ TCID_50_.ml^-1^) within 12-24 h pi (Fig. 1B). Interestingly, while SARS-CoV-2 also replicated to high titers in DL cells (∼10^6^ to 10^7^ TCID_50_.ml^-1^), replication kinetics was delayed with peak viral titers being reached by 72 h pi (Fig. 1B). These results show that DL cells are susceptible to SARS-CoV-2 infection and support productive virus replication *in vitro*.

### Infection and transmission of SARS-CoV-2 in white-tailed-deer

Given that white-tailed-deer have been recently identified as one of the species with an ACE2 protein with high binding probability to the SARS-CoV-2 S protein (18), we investigated the susceptibility of white-tailed-deer fawns to SARS-CoV-2 infection. Six-week-old fawns (n = 4) were inoculated intranasally with 5 ml (2.5 ml per nostril) of a virus suspension containing 10^6.3^ TCID_50_.ml^-1^ of SARS-CoV-2 TGR/NY/20, an animal SARS-CoV-2 strain that is identical to human viral strains (22). To assess the potential transmission of SARS-CoV-2 between white-tailed-deer, two fawns (n = 2) were maintained as non-inoculated contacts in the same biosafety level 3 (BSL-3Ag) room. The inoculated and indirect contact animals were kept in separate pens divided by a plexiglass barrier, to prevent direct nose-to-nose contact between inoculated and contact animals (Fig. 2A). Following inoculation, animals were monitored daily for clinical signs and body temperature. No clinical signs or overt respiratory distress were observed in any of the inoculated or contact animals during the 21-day experimental period. Interestingly, a slight and transient increase in body temperature was noted in 3/4 (no. 2001, 2042 and 2043) inoculated fawns between days 1-3 pi (Fig. 2B). One fawn (no. 2001) died on day 8 pi of unrelated intestinal perforation and peritonitis. The body temperature in both indirect contact animals (no. 2006 and 2044) remained within physiological ranges throughout the experimental period (Fig. 2B). Postmortem examination of fawns at 8 (animal no. 2001) or 21 days pi (animals no. 2042, 2043, 2045, 2005 and 2044) revealed no gross lesions. Microscopic changes observed were not associated with the presence of the virus, as no viral RNA was detected by PCR or ISH (Suppl data).

**Figure 2.**
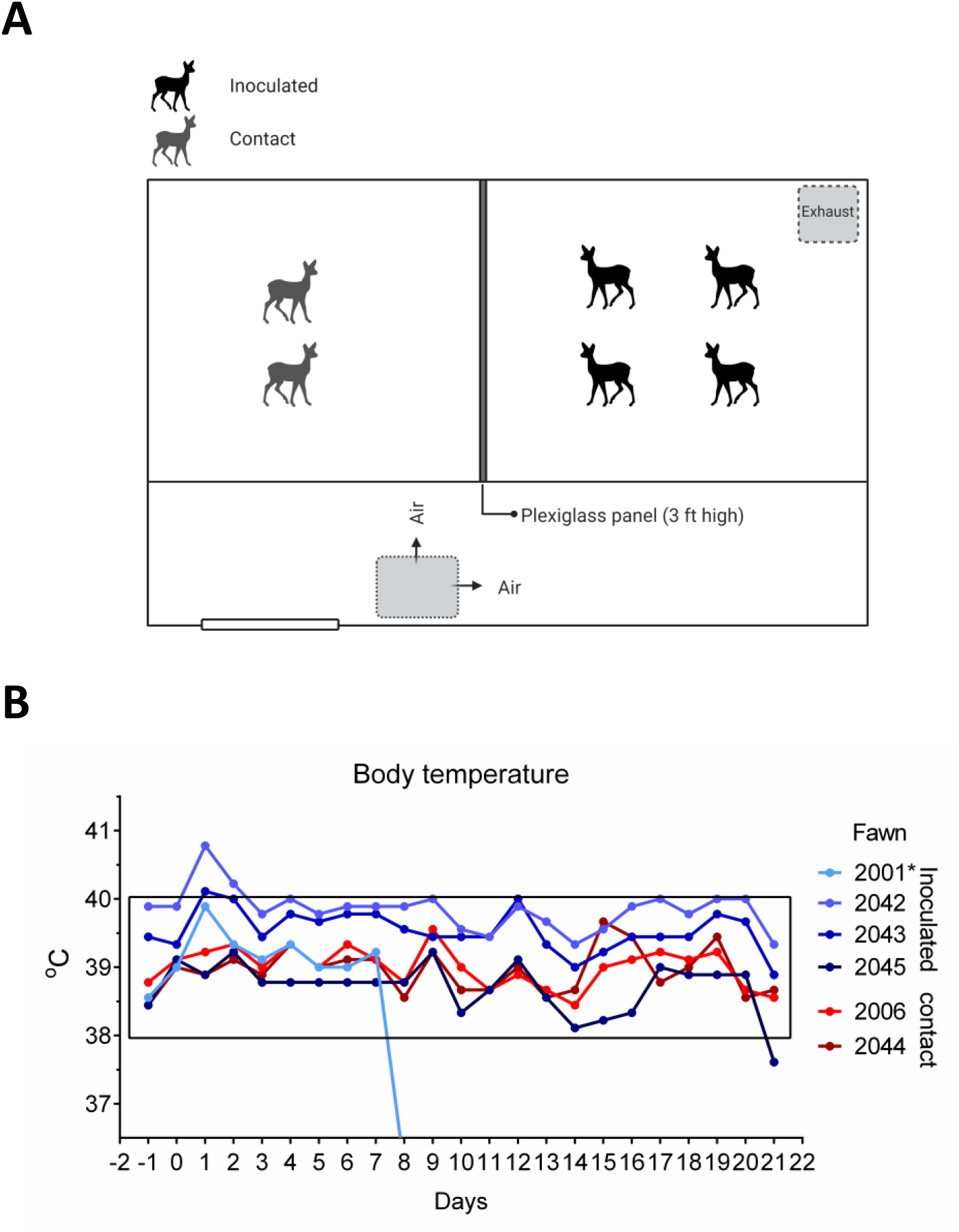
Infection and transmission of SARS-CoV-2 in white-tailed-deer. (A) Room set up of animal experiment. Fawns were kept in a room of biosafety level 3 (Agriculture) (BSL-3Ag) facility. Four fawns were inoculated intranasally with a virus suspension containing 5×10^6.3^ TCID_50_ of SARS-CoV-2 isolate TGR/NY/20, and two fawns were maintained as non-inoculated room contact animals. All fawns were maintained in a 3.7m x 3.7m room, and inoculated and room contact animals were kept in two pens separated by a plexiglass barrier approximately 0.9 m (∼3-feet) in height, to prevent direct nose-to-nose contact. Airflow in the room was maintained at 10-11 air exchanges per hour and was directional from the contact pen towards the inoculated pen. (B) Fawns were microchipped subcutaneously for identification and monitored daily for clinical signs and body temperature starting on day 1 before inoculation or contact day (day −1). Body temperatures are expressed in degrees Celsius (°C).

Replication of SARS-CoV-2 in the upper respiratory and gastrointestinal tracts and the dynamics and patterns of virus shedding and viremia were assessed in inoculated and indirect contact fawns. Nasal secretions and feces were collected by nasal and rectal swabs on days 0, 1, 2, 3, 4, 5, 6, 7, 10, 12, 14 and 21 pi, serum and buffy coat collected on days 0, 7, 14, and 21 pi were subjected to nucleic acid extraction and tested for the presence of SARS-CoV-2 RNA by real-time reverse transcriptase PCR (RT-rPCR). While viremia was not detected in serum and buffy coat samples, viral RNA was detected between days 2 and 21 pi in nasal secretions from inoculated animals (Fig. 3A), with higher viral RNA loads being detected between days 2 and 7 pi and decreasing thereafter through day 21 pi (Fig. 3A). Notably, high levels of viral RNA were also detected in nasal secretions from indirect contact animals throughout the experimental period (Fig. 3B). Viral RNA from the feces was detected in all inoculated and indirect contact animals (Fig. 3A and B), however, intermittent and short duration fecal shedding was observed, with most animals (4/5) only transiently shedding detectable SARS-CoV-2 in feces through days 6-7 pi (Fig. 3A and B).

**Figure 3.**
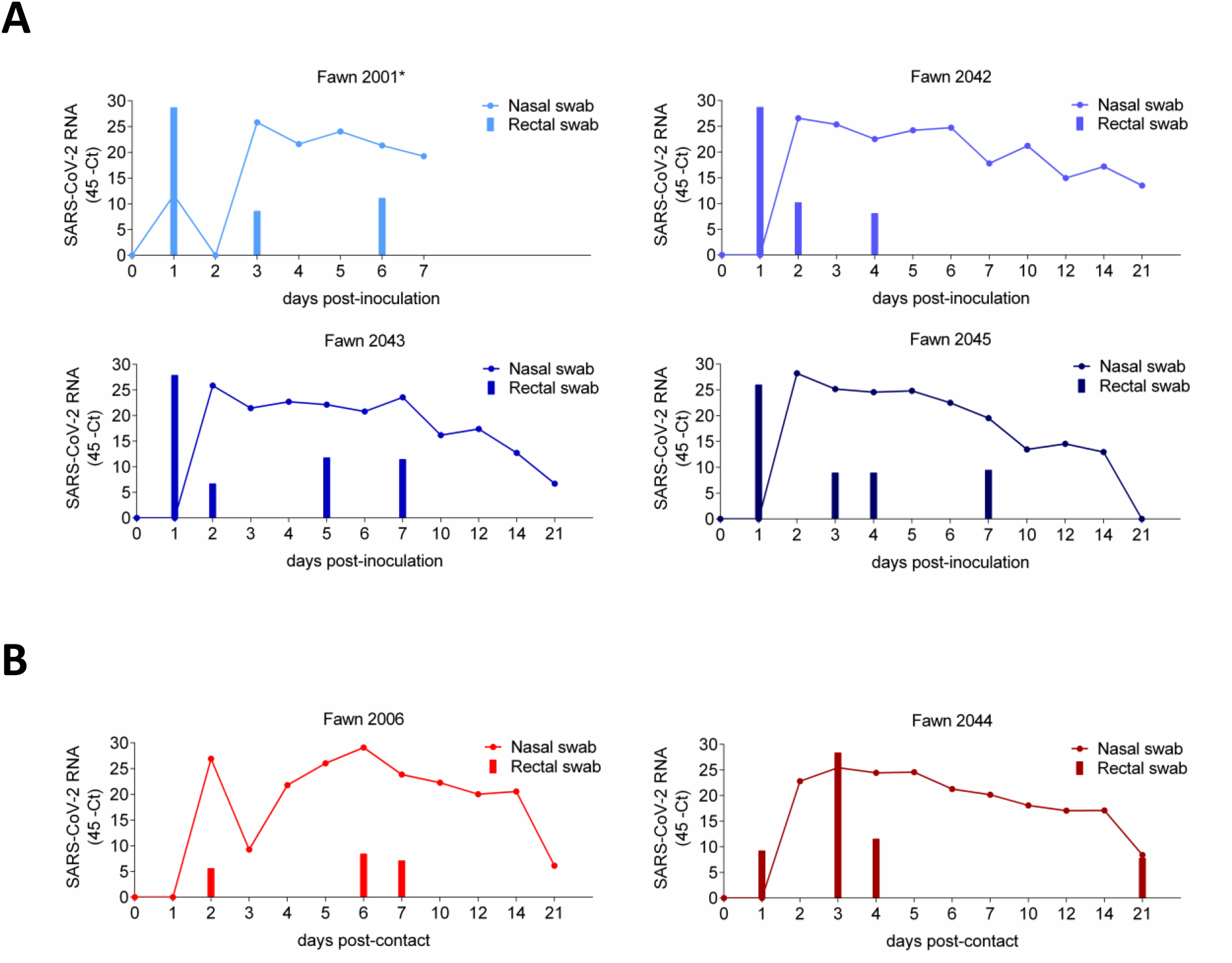
Viral RNA in nasal secretion and feces. (A) The dynamics of virus shedding was assessed in nasal secretions (line) and feces (bars) of four inoculated fawns (no 2001, 2042, 2043, and 2045). Nasal and rectal swabs collected on days 0-7, 10, 12, 14 and 21 post-inoculation (pi), were tested for the presence of SARS-CoV-2 RNA by real-time reverse transcriptase PCR (RT-rPCR). (B) The dynamics of virus shedding of contact fawns (no 2001 and 2044). Nasal and rectal swabs collected on the same time-points as the inoculated animals, were subjected to nucleic acid extraction and tested for the presence of SARS-CoV-2 RNA by RT-rPCR.

Notably, infectious SARS-CoV-2 was detected in nasal secretions of all inoculated and indirect contact animals between days 2 and 5 pi (Fig. 4A). Indirect contact animals shed infectious SARS-CoV-2 in nasal secretions through day 7 pi. Viral titers shed in nasal secretions ranged from 2.0 to 4.8 log_10_ TCID_50_.ml^-1^ (Fig. 4A). Infectious SARS-CoV-2 shedding in feces was detected in inoculated animals on day 1 pi (Fig. 4B). Sequencing of SARS-CoV-2 from select nasal secretions from inoculated and indirect contact animals confirmed the identity of the virus, matching the sequence of the inoculated virus. Together these results indicate that SARS-CoV-2 productively infected and replicated in the upper respiratory tract of white-tailed-deer. In addition, the virus was efficiently transmitted to indirect contact animals.

**Figure 4.**
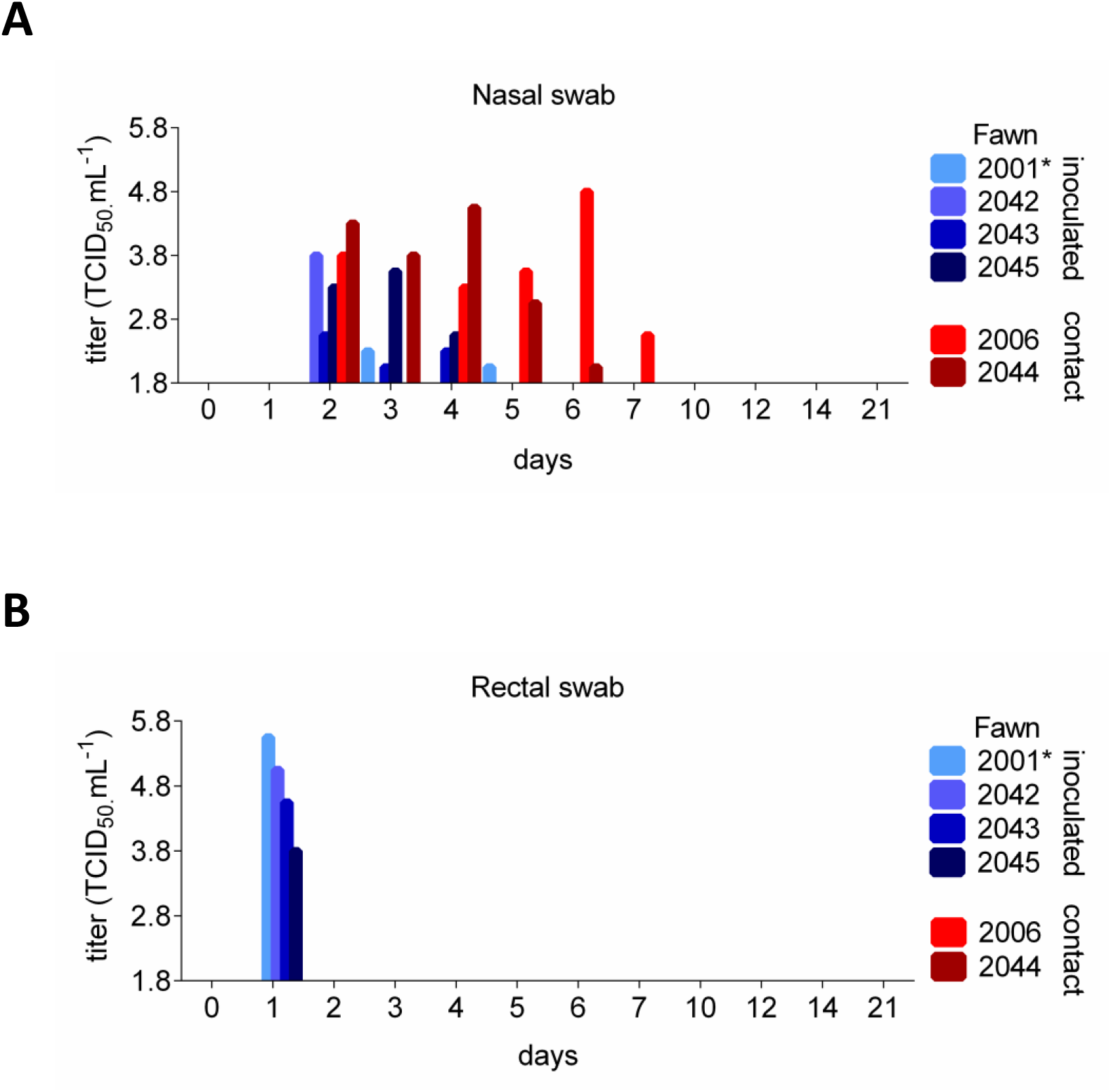
Shedding of infectious SARS-CoV-2 by inoculated and contact fawns. (A) Infectious virus was assessed by virus isolation in nasal secretions in RT-rPCR-positive samples. (B) Infectious SARS-CoV-2 shedding in feces in RT-rPCR-positive samples. Virus titers were determined using end point dilutions and the Spearman and Karber’s method and expressed as TCID_50_.ml^-1^.

### Viral load and tissue distribution

Viral RNA and tissue distribution of SARS-CoV-2 were assessed on day 8 (animal no. 2001, which died of unrelated cause [intestinal perforation]) and 21 pi (animals no. 2042, 2043, 2045, 2005 and 2044). Tissues were collected and processed for RT-rPCR, virus isolation and *in situ* hybridization (ISH). On day 8 pi (fawn 2001), SARS-CoV-2 RNA was detected in nasal turbinates, palatine tonsil, spleen, ileocecal junction and in submandibular-, medial retropharyngeal-, tracheobronchial- and mediastinal lymph nodes (Fig.5). Among the tissues collected on day 21 pi, from the remaining inoculated and indirect contact fawns, SARS-CoV-2 RNA was consistently detected in nasal turbinates, palatine tonsil and retropharyngeal lymph node (Fig. 5). This subset of tissues presented the higher viral RNA loads on both days 8 and 21 pi, when compared to the other tissues tested (Fig. 5).

**Figure 5.**
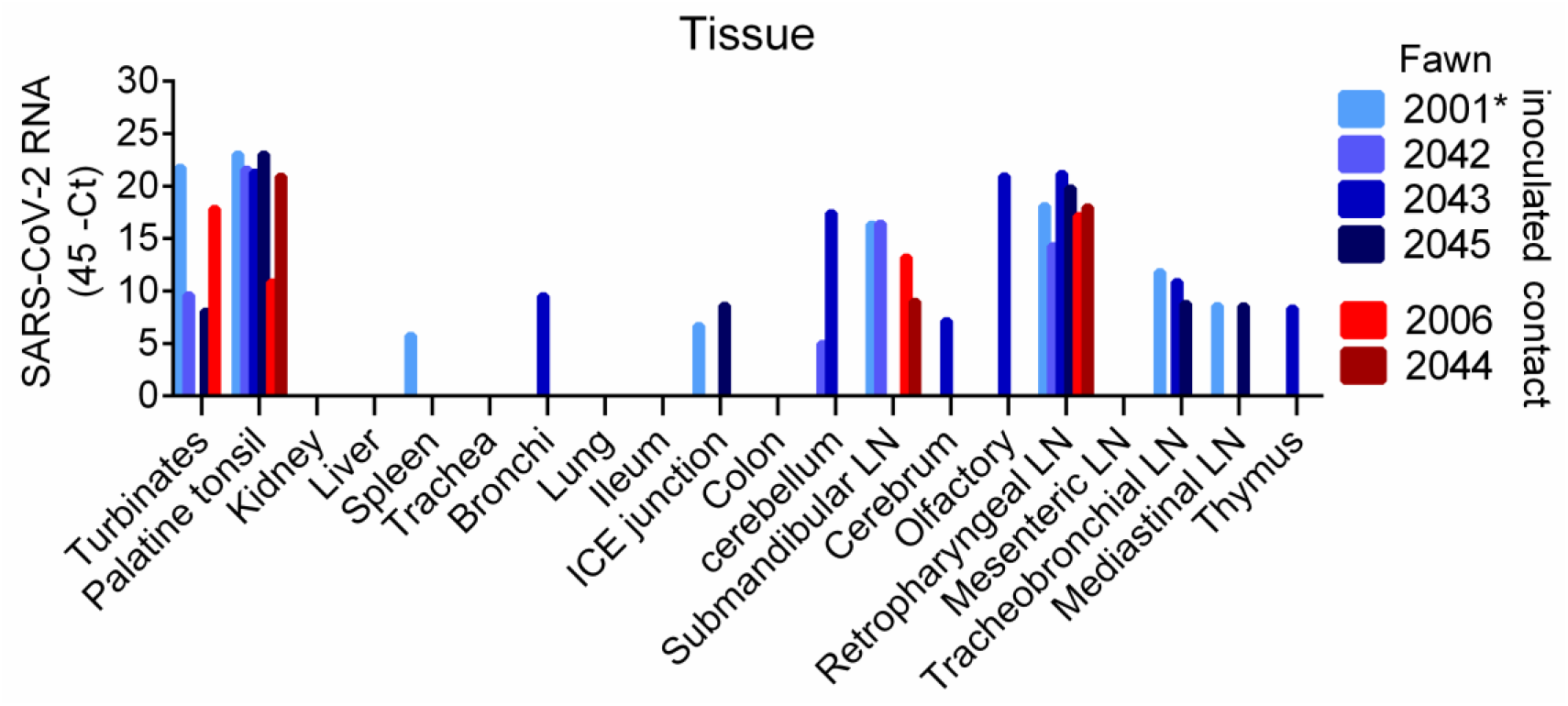
Tissue distribution of SARS-CoV-2 RNA. Tissues were collected and processed for RT-rPCR on day 8 post-inoculation (pi) (animal no. 2001, which died of unrelated cause) and 21 pi of the remaining animals (animals no. 2042, 2043, 2045, 2005 and 2044).

The presence of SARS-CoV-2 RNA was confirmed by ISH, in the palatine tonsils and medial retropharyngeal lymph nodes of all inoculated and indirect contact fawns. Labeling of viral RNA was intense and limited to central regions of lymphoid follicles (Fig. 6A, B). Viral RNA was also noted in tracheobronchial and/or mediastinal lymph nodes of 5/6 deer, but was less intense than that seen in tonsils, and was limited to lymph node medulla (Fig. 6C). Only the nasal turbinates of fawn 2001, collected on day 8 pi exhibited viral RNA labeling. Within the nasal lumen there was strong labeling of aggregates of mucus and cell debris (Fig. 6D). No viral RNA was detected in sections of lung, kidney, brain, intestine, or mesenteric lymph nodes.

**Figure 6.**
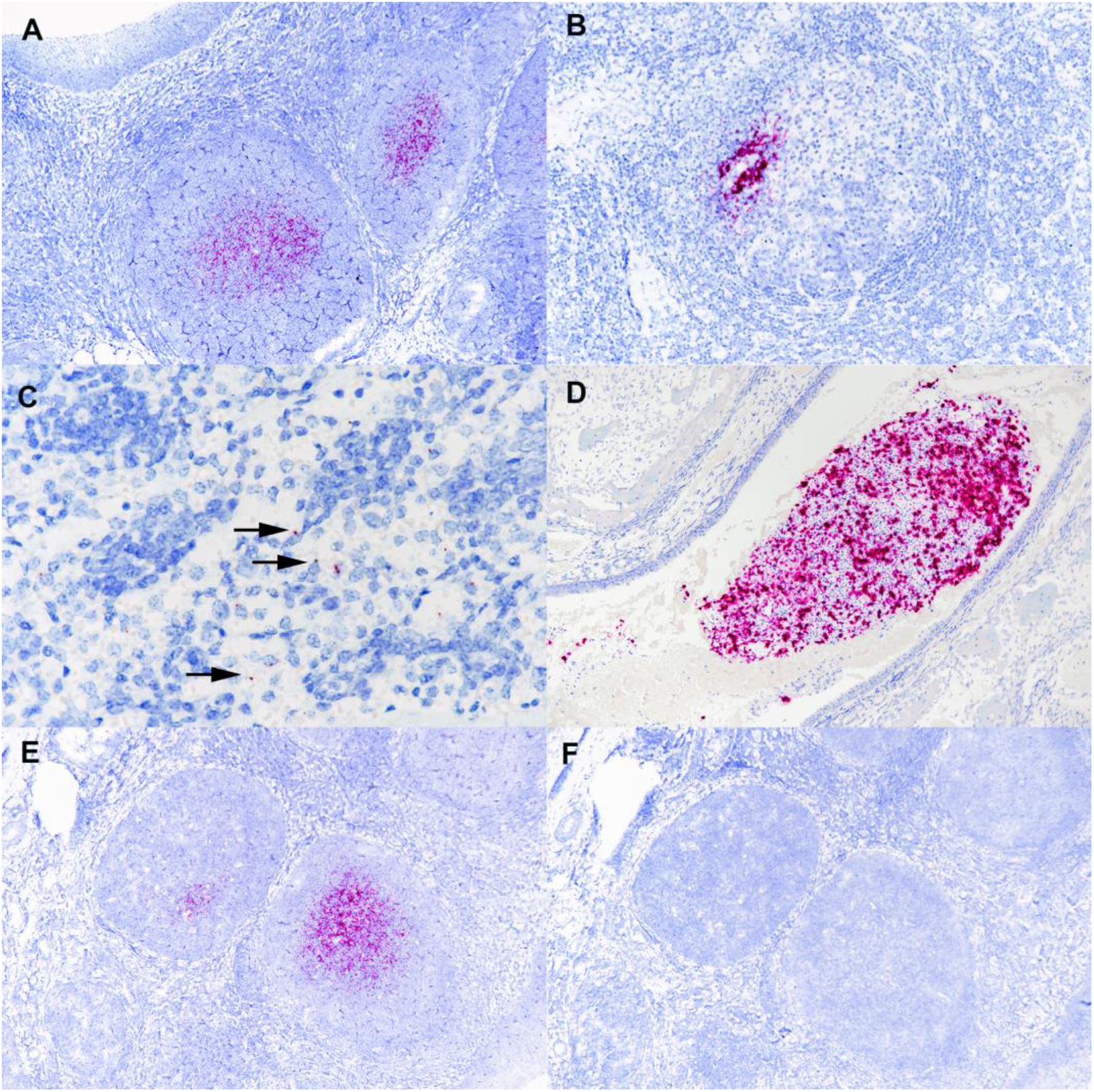
Tissues from white-tailed deer fawn inoculated intranasally with SARS-CoV-2 and examined 21 days later. Note intense labeling of viral RNA in the centers of lymphoid follicles (A) located subjacent to tonsillar epithelium (upper left). Note labeling for SARS-CoV-2 RNA within medial retropharyngeal lymph node follicle (B) and mediastinal lymph node medulla (C). Nasal turbinate lumen contains aggregate of mucus, cells and debris with intense labeling for SARS-CoV-2 RNA (D). Adjacent microscopic sections demonstrate intense labeling of lymphoid follicles with probe for SARS-CoV-2 RNA (E) but no labeling using the anti-genomic sense probe (F). ISH-RNAscope.

Tissues were also analyzed using the anti-genomic negative sense RNA probe (V-nCoV2019-S-sense probe) to determine virus replication. While intense labeling was noted with the genomic probe (Fig 6E), no labeling was observed with the anti-sense probe in any of the tissues (Fig.6F), suggesting lack of virus replication. These findings are consistent with lack of virus isolation from RT-rPCR positive tissues. The ability of the anti-sense probe to bind actively replicating virus, was confirmed in Vero cells infected with SARS-CoV-2 isolate TGR/NY/20 (Suppl Fig. 1).

### Antibody responses in white-tailed deer following SARS-CoV-2 infection

The serological responses to SARS-CoV-2 were assessed using a Luminex and a virus neutralization (VN) assay. Serum samples collected on days 0, 7, 14 and 21 pi were used to assess nucleocapsid (N)- and S-receptor binding domain (RBD)-specific or neutralizing antibodies (NA) levels following SARS-CoV-2 infection. All inoculated and indirect contact animals seroconverted to SARS-CoV-2, with antibodies against S-RBD and NA being detected as early as day 7 pi (Fig. 7B and 7C, respectively) and increasing thereafter on days 14 and 21 pi. Overall, levels of antibodies against the N-protein were lower and appeared at later time points (day 14-21) (Fig. 7A). Geometric mean titers (GMT) of NA ranged from 37 to 107, 85 to 214, and 85 to 256 on days 7, 14, and 21, respectively (Fig. 7C). These results confirm infection of all inoculated fawns and demonstrate transmission to indirect contact animals.

**Figure 7.**
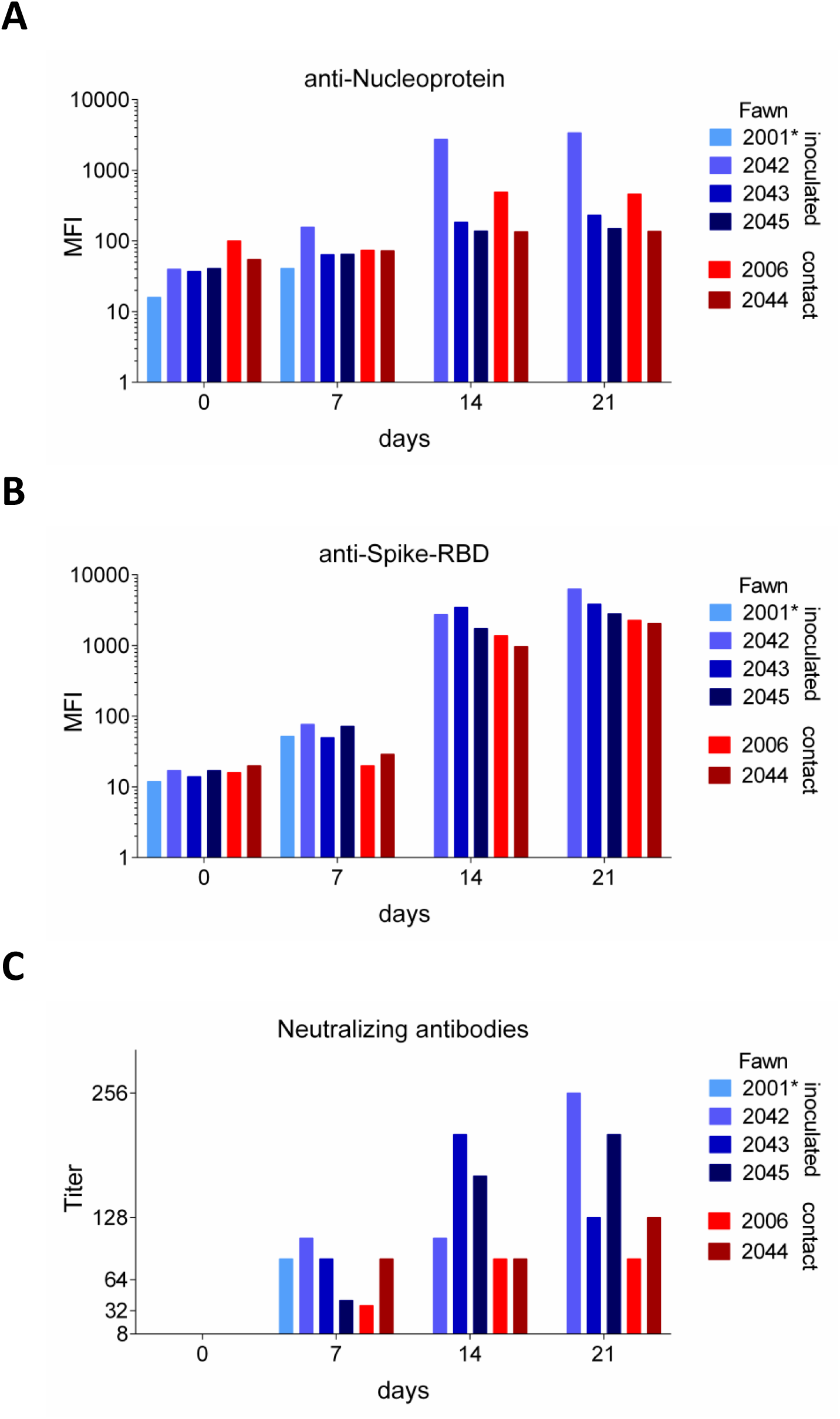
Antibody responses in white-tailed deer following SARS-CoV-2 infection. (A) Luminex assay to assess IgG anti-SARS-CoV-2-nucleocapsid (N) in serum samples collected on days 0, 7, 14 and 21 post-inoculation (pi) (fawns no 2001, 2042, 2043, and 2045), or contact (fawns no 2006 and 2044). (B) Luminex assay to assess IgG anti-SARS-CoV-2-S-receptor binding domain (RBD)-specific at the same time point and fawns as above. (C) Virus neutralization (VN) assay. Neutralizing antibody (NA) titers were expressed as the reciprocal of the highest dilution of serum that completely inhibited SARS-CoV-2 infection/replication in serum at the same time point and fawns as above. Result represent the geometric mean titers (GMT) of three independent experiments.

## Discussion

Here we confirmed *in silico* predictions (18), demonstrating that white-tailed deer are susceptible to SARS-CoV-2 infection. Infection of deer cells *in vitro* resulted in productive virus replication. Most importantly, intranasal inoculation of white-tailed-deer fawns led to subclinical infection and productive virus replication in the upper respiratory tract with shedding of infectious virus in nasal secretions of infected animals.

One of the most remarkable characteristics of SARS-CoV-2 is its highly efficient transmissibility (35, 36). Consistent with this, rapid SARS-CoV-2 transmission was observed in several experimentally infected animal species (27, 31, 37–43). Similarly, results here show efficient SARS-CoV-2 transmission between white-tailed-deer. Indirect contact fawns became infected and supported productive SARS-CoV-2 replication as evidenced by virus shedding in nasal secretions and seroconversion. Evidence to date indicates that transmission of SARS-CoV-2 occurs mainly through direct, indirect, or close contact with infected individuals through infected secretions such as respiratory-, salivary-, and/or fecal droplets (44). Under experimental conditions, transmission via direct contact has been demonstrated in ferrets, minks, raccoon dogs, hamsters, cats, and deer mice (28, 31, 37, 39, 40, 43, 45), while indirect/aerosol transmission was observed in ferrets and hamsters (37, 39). The experimental setup in the present study prevented direct contact between animals suggesting that transmission of SARS-CoV-2 from inoculated to room indirect contact animals most likely occurred via aerosols or droplets generated by inoculated animals.

Despite the potential for severe illness and mortality due to SARS-CoV-2 infection in humans (46, 47), asymptomatic and mild cases represent approximately 80% of human COVID-19 cases, while severe and critical disease outcomes account for approximately 15 and 5% of the cases, respectively (48). Notably, experimental infections in several animal species, including non-human primates (26), cats (19, 23, 49), ferrets (23, 31, 50, 51), minks (27), deer mice (40), and racoon dogs (42) have mostly resulted in subclinical infections or only mild respiratory distress. Consistent with these observations, white-tailed-deer fawns inoculated here did not present overt clinical disease following SARS-CoV-2 infection. Only a transient and modest increase in body temperature was observed in 3 of 4 inoculated animals between days 1-3 pi. It is important to note that most severe and critical cases of COVID-19 in humans have been associated with underlying clinical conditions (e.g. heart and pulmonary disease, cancer, and diabetes, among others) which may complicate full recapitulation of the clinical manifestations of the human disease in animal species. Differences in expression and/or tissue distribution of the SARS-CoV-2 receptor (ACE2) (Suppl. Fig. 2), of cellular proteases (transmembrane serine protease 2, TMPRRS2, cathepsins) or other co-factor(s) that might be required for efficient virus infection and replication in target tissues, may help to explain the diverse clinical outcomes of infection in humans and animals.

While no viremia was detected in infected animals, active virus shedding was observed in nasal secretions of both inoculated and indirect contact animals. The patterns and dynamics of SARS-CoV-2 shedding observed in fawns here are consistent with what has been described for other susceptible animal species (23, 31, 40, 42, 49, 52, 53). Most importantly, the duration and magnitude of virus shedding observed in deer corroborate observations in humans, in which prolonged viral RNA has been detected by RT-rPCR (21-35 days), with virus infectivity being usually limited to the first 7-10 days of infection (54, 55). This could be partially explained by the activity of effector host immune responses elicited against the virus, especially of neutralizing antibodies, which were detected as early as day 7 pi in all fawns here, paralleling decreased virus infectivity in nasal secretions.

Viral RNA load and tissue distribution were assessed on day 8 pi (animal no. 2001) following acute infection and on day 21 pi after termination of the experiment (remaining fawns). High viral RNA loads were detected in nasal turbinates, palatine tonsils, and medial retropharyngeal lymph nodes, suggesting that these tissues may potentially serve as sites of SARS-CoV-2 replication in white-tailed-deer. Since no infectious virus was recovered from any of the tissues sampled on days 8 or 21 pi, additional studies involving serial tissue collection over time are needed to determine specific sites of virus replication. Interestingly, no viral RNA was detected in the lung tissues from inoculated or indirect contact fawns, suggesting that these tissues may not be targeted by SARS-CoV-2 following infection of white-tailed-deer. The possibility that the virus may have been cleared from these sites by the time of tissue collection, however, cannot be excluded.

Microscopic changes were observed in the lungs of 2 of 4 inoculated fawns and the 1 indirect contact fawn examined 21 days pi. These changes were characterized by congestion, lymphohistiocytic interstitial pneumonia, hyaline membranes, and intra-alveolar fibrin, all of which are characteristic of the acute phase of diffuse alveolar damage, the predominant finding in the lungs of patients affected with COVID-19 (60, 61) (Suppl Fig. 3). Mild to moderate pulmonary changes have been described in other species following experimental SARS-CoV-2 infection, including domestic cats (41), rhesus macaques (25, 56, 57), hACE2 mice (58, 59), ferrets (37), and minks (27). It is also notable that no SARS-CoV-2 viral RNA was associated with lesions in these white-tailed deer, which could be a result of earlier viral clearance from these sites, or potent immunological responses following SARS-CoV-2 infection.

In summary, our study shows that white-tailed-deer are susceptible to SARS-CoV-2 infection and can transmit the virus to indirect contact animals. These results confirmed *in silico* predictions describing a high propensity of interaction between SARS-CoV-2 S protein and the cervid ACE2 receptor. Our findings indicate that deer and other cervids should be considered in investigations conducted to identify the origin and potential intermediate host species that may have served as the link host reservoir to humans.

## Materials and Methods

### Cells and Virus

Vero cells (ATCC^®^ CCL-81^™^), Vero E6 (ATCC^®^ CRL-1586^™^), and Vero E6/TMPRSS2 (JCRB Cell Bank, JCRB1819) were cultured in Dulbecco’s modified eagle medium (DMEM), while deer lung (DL) cells were cultured in minimum essential medium (MEM). Both DMEM and MEM were supplemented with 10% fetal bovine serum (FBS), L-glutamine (2mM), penicillin (100 U.ml^−1^), streptomycin (100 μg.ml^−1^) and gentamycin (50 μg.ml^−1^). The cell cultures were maintained at 37 °C with 5% CO_2_. The SARS-CoV-2 isolate TGR/NY/20 obtained from a Malayan tiger naturally infected with SARS-CoV-2 and presenting with respiratory disease compatible with SARS-CoV-2 infection (22) was propagated in Vero CCL-81 cells. Low passage virus stocks (passage 4) were prepared, cleared by centrifugation (1966 x *g* for 10 min) and stored at −80 °C. The endpoint titer was determined by limiting dilution following the Spearman and Karber method. A viral suspension containing 10^6.3^ tissue culture infectious dose 50 per ml (TCID_50_.ml^-1^) was used for all *in vitro* experiments and fawn inoculations.

### Cell susceptibility and growth curves

The susceptibility and kinetics of replication of the SARS-CoV-2 in DL cells was assessed *in vitro* and compared to virus replication in Vero E6 and Vero E6/TMPRSS2. For this, DL, Vero E6 and Vero E6/TMPRSS2 cells were inoculated with SARS-CoV-2 isolate TGR/NY/20 at a multiplicity of infection of 1 (MOI = 1) and the cells incubated for 24 h at 37 °C with 5% CO_2_. At 24 h post-inoculation, cells were fixed with 3.7% formaldehyde for 30 min at room temperature, permeabilized with 0.2% Triton X-100 (in Phosphate Buffered Saline [PBS]) and subjected to an immunofluorescence assay (IFA) using a monoclonal antibody (MAb) anti-ACE2 (Sigma-Aldrich), and then incubated with a goat anti-rabbit IgG (goat anti-rabbit IgG, Alexa Fluor 488^®^), and using a monoclonal antibody (MAb) anti-SARS-CoV-2 nucleoprotein (N) (clone B6G11) produced and characterized in Dr. Diel’s laboratory, and then incubated with a goat anti-mouse IgG secondary antibody (goat anti-mouse IgG, Alexa Fluor^®^ 594), and Nuclear counterstain was performed with DAPI, and visualized under a fluorescence microscope. To assess the kinetics of replication of SARS-CoV-2 isolate TGR/NY/20, Vero E6, Vero E6/TMPRSS2 and DL cells were cultured in 12-well plates, inoculated with SARS-Cov-2 isolate TGR/NY/20 (MOI = 0.1 and MOI = 1 and harvested at various time points post-inoculation (12, 24, 36, 48 and 72 h pi). Virus titers were determined on each time point using end-point dilutions and the Spearman and Karber’s method and expressed as TCID_50_.ml^-1^.

### Animals, inoculation and sampling

All animals were handled in accordance with the Animal Welfare Act Amendments (7 U.S. Code §2131 to §2156) and all study procedures were reviewed and approved by the Institutional Animal Care and Use Committee at the National Animal Disease Center (IACUC approval number ARS-2020-861). White-tailed deer fawns (n = 6) were obtained from a breeding herd maintained at the National Animal Disease Center in Ames, IA. Within 24 hrs of birth, fawns were removed from the pasture, moved to indoor housing, and bottle-fed. Hand-raising fawns has been found to greatly decrease animal stress when moved to containment housing (62). At approximately 4 weeks of age, fawns were moved to the biosafety level 3 (Agriculture) (BSL-3Ag) facility at NADC, and were microchipped (subcutaneously; SC) for identification and body temperature monitoring. Fawns were fed white-tailed deer milk replacer and hay was also available.

After a 2-week acclimation period, four fawns were inoculated intranasally with 5 ml (2.5 ml per nostril) of a virus suspension containing 10^6.3^ TCID_50_.ml^-1^ of SARS-CoV-2, isolated from respiratory secretions from a tiger at the Bronx Zoo (TGR/NY/20) (32). Two fawns were maintained as non-inoculated indirect contact animals to evaluate potential transmission of SARS-CoV-2 from inoculated to indirect contact animals. All fawns were maintained in a 3.7m x 3.7m room, and inoculated and room indirect contact animals were kept in two pens separated by a plexiglass barrier approximately 0.9 m (∼3-feet) in height, to prevent direct nose-to-nose contact (Fig. 2A). Airflow in the room was maintained at 10-11 air exchanges per hour, at a standard exchange rate for BSL-3Ag housing of large animals. Body temperatures were recorded daily, nasal swabs (NS) and rectal swabs (RS) were collected on days 0, 1, 2, 3, 4, 5, 6, 7, 10, 12, 14 and 21 post-inoculation (pi). Upon collection, swabs were placed individually in sterile tubes containing 2 ml of viral transport media (MEM with 1,000 U.ml^-1^ of penicillin, 1,000 µg.ml^-1^ of streptomycin, and 2.5 µg.ml^-1^ of amphotericin B) and stored at −80 °C until analysis. Blood was collected through jugular venipuncture in EDTA and serum separator tubes on days 0, 7, 14, and 21 pi. The tubes were centrifuged for 25 min at 1200 x *g* and buffy coat (BC) was collected from EDTA tubes and stored at −80 °C. Serum from the serum separator tubes was aliquoted and stored at −80 °C until analysis.

Fawns were humanely euthanized on day 21 pi. Following necropsy, multiple specimens including tracheal wash (TW), lung lavage (LL) and several tissues (nasal turbinates, palatine tonsil, thymus, trachea, lung, bronchi, kidney, liver, spleen, ileum, ileocecal junction, spiral colon, cerebellum, cerebrum, olfactory bulbs and medial retropharyngeal, mandibular, tracheobronchial, mediastinal and mesenteric lymph nodes) were collected. Samples were processed for RT-rPCR and virus isolation (VI) and were individually bagged, placed on dry ice, and transferred to a −80 °C freezer until testing. Additionally, tissue samples were collected and processed for standard microscopic examination and *in situ* hybridization (ISH). For this, tissue fragments of approximately ≤0.5 cm in width were fixed by immersion in 10% neutral buffered formalin (≥20 volumes fixative to 1 volume tissue) for approximately 24 h, and then transferred to 70% ethanol, followed by standard paraffin embedding techniques. Slides for standard microscopic examination were stained with hematoxylin and eosin (HE).

### Nucleic acid extraction and real-time RT-PCR (RT-rPCR)

Nucleic acid was extracted from nasal secretions, feces, BC, serum tracheal wash, lung lavage and all the tissue samples collected at necropsy. Before extraction, 0.5 g of tissues were minced with a sterile scalpel and resuspended in 5 ml DMEM (10% w/v) and homogenized using a stomacher (Stomacher^®^ 80 Biomaster; one speed cycle of 60s). Homogenized tissue samples were cleared by centrifugation (1966 x *g* for 10 min) and 200 µL of the homogenate supernatant used for RNA extraction using the MagMax Core extraction kit (Thermo Fisher, Waltham, MA, USA) and the automated KingFisher Flex nucleic acid extractor (Thermo Fisher) following the manufacturer’s recommendations. The real-time reverse transcriptase PCR (RT-rPCR) was performed using the EZ-SARS-CoV-2 Real-Time RT-PCR assay (Tetracore Inc., Rockville, MD). An internal inhibition control was included in all reactions. Positive and negative amplification controls were run side-by-side with test samples. All RNA extractions and RT-rPCR were performed at the Cornell Animal Health Diagnostic Center (AHDC).

### Virus Isolation and titration

Nasal and rectal swabs collected on days 0, 1, 2, 3, 4, 5, 6, 7, 10, 12, 14 and 21, and tissues collected during the necropsy that tested positive for SARS-CoV-2 by RT-rPCR were subjected to virus isolation under biosafety level 3 conditions at Cornell University. Twenty-four well plates were seeded with ∼75,000 Vero E6/TMPRSS2 cells per well 24 h prior to sample inoculation. Cells were rinsed with phosphate buffered saline (PBS) (Corning^®^) and inoculated with 150 µl of each sample and inoculum adsorbed for 1 h at 37 °C with 5% CO_2_. Mock-inoculated cells were used as negative controls. After adsorption, replacement cell culture media supplemented as described above was added, and cells were incubated at 37 °C with 5% CO_2_ and monitored daily for cytopathic effect (CPE) for 3 days. SARS-CoV-2 infection in CPE-positive cultures was confirmed with an immunofluorescence assay (IFA) as described above. Cell cultures with no CPE were frozen, thawed, and subjected to two additional blind passages/inoculations in Vero E6/TMPRSS2 cell cultures. At the end of the third passage, the cells cultures were subjected to IFA as above. Positive samples were subjected to end point titrations by limiting dilution using the Vero E6/TMPRSS2 cells and virus titers were determined using the Spearman and Karber’s method and expressed as TCID_50_.ml^-1^.

### In situ hybridization (ISH)

Paraffin-embedded tissues were sectioned at 5 µm and subjected to ISH using the RNAscope ZZ probe technology (Advanced Cell Diagnostics, Newark, CA). *In situ* hybridization was performed to detect tissue distribution of SARS-CoV-2 nucleic acid in palatine tonsil, medial retropharyngeal-tracheobronchial-mediastinal and mesenteric lymph nodes, nasal turbinate, brain, lung, and kidney, using the RNAscope 2.5 HD Reagents–RED kit (Advanced Cell Diagnostics) as previously described (63). Proprietary ZZ probes targeting SARS-CoV-2 RNA (V-nCoV2019-S probe ref# 8485561) or anti-genomic RNA (V-nCoV2019-S-sense ref#845701) designed and manufactured by Advance Cell Diagnostics were used for detection of viral RNA. A positive control probe targeted the *Bos taurus* –specific cyclophilin B (PPIB Cat# 3194510) or ubiquitin (UBC Cat # 464851) housekeeping genes, while a probe targeting dapB of *Bacillus subtilis* (Cat # 312038) was used as a negative control. The distribution of mRNA of ACE2 was assessed using the BaseScope 2.5 HD Reagents–RED kit (Advanced Cell Diagnostics) as previously described (63). Proprietary ZZ probes targeting the region spanning AA 31-82 for the ACE2 receptor specific to *Odocoileus virginianus* (BA-Ov-ACE2-1zz-st probe; ref# 900101) designed and manufactured by Advance Cell Diagnostics. Slides were counterstained with hematoxylin, and examined by light microscopy using a Nikon Eclipse Ci microscope. Digital images were captured using a Nikon DE-Ri2 camera.

### Serological analysis

Antibody responses to SARS-CoV-2 was assessed by a Luminex and virus neutralization assays developed *in house*. For the Luminex assay, SARS-CoV-2 antigens were expressed as IL-4 fusion proteins in mammalian cells as previously described (64). SARS-CoV-2 RNA (strain Hu-WA-1) was derived from Vero cells infected with the virus and cDNA was synthesized using the SuperScript III Reverse Transcriptase (Life Technologies) and oligo dT and six hexamer random primers. The cDNA was used to amplify the receptor binding domain (amino acids 319 to 529; RBD) of the Spike protein and the whole nucleocapsid protein (amino acids 1 – 419; NP) by PCR. The primer sequences for RBD were AA<u>GGATCC</u>AAGAGTCCAACCAACAGAATCTATTGTT (forward) and AAA<u>GGTACC</u>TTACTTTTTAGGTCCACAAACAGTTGCT (reverse). NP primers were AA<u>GGATCC</u>AATGTCTGATAATGGACCCCAAAATC (forward) and AAA<u>GGTACC</u>TTAGGCCTGAGTTGAGTCAGCACTG (reverse). The restriction sites for expression cloning are underlined. Both PCR products were cloned into the multiple cloning site of the mammalian expression vector pcDNA3.1 (ThermoFisher Scientific, Waltham, MA, USA) downstream of the equine IL-4 sequence, as described previously (64). Nucleotide sequences of RBD and NP were verified by Sanger sequencing and identical to respective sequences of the Hu-WA-1 strain (GenBank accession number NC_045512).

CHO-K1 cells were transiently transfected with the recombinant plasmid constructs. Expression and secretion of the recombinant SARS-CoV-2 antigen fusion proteins by CHO transfectants was confirmed using the IL-4 tag by flow cytometric analysis and ELISA, as previously described (64). After 24-30 hrs of incubation, the supernatants were harvested for fluorescent bead-based multiplex assays.

Fluorescent beads were coupled with the anti-equine IL-4 antibody, clone 25 (RRID: AB_2737308) as previously described (65). The RBD/IL-4 and NP/IL-4 fusion proteins were bound by incubating the anti-IL-4 coupled beads with the fusion protein supernatant solution for 30 min at room temperature, followed by a wash. The assay was performed with a few modifications from previously described procedures (65, 66). Briefly, beads were incubated with deer serum samples diluted 1:100. The assay was detected using a biotinylated mouse anti-goat IgG (H+L) (RRID: AB_2339061, Jackson Immunoresearch Laboratories, West Grove, PA) cross-reactive with deer immunoglobulin. Afterwards, streptavidin-phycoerythrin (Invitrogen, Carlsbad, CA) was added as a final detection step. All incubation steps were for 30 min at room temperature and the assay was washed after each incubation step. The assay was measured in a Luminex 200 instrument (Luminex Corp., Austin, TX). Assay results were expressed as median fluorescence intensity (MFI).

Neutralizing antibody responses to SARS-CoV-2 was assessed by a virus neutralization assay (VNA) performed under BSL-3 conditions at the Cornell AHDC. Twofold serial dilutions (1:8 to 1:4,096) of serum samples were incubated with 100 − 200 TCID_50_ of SARS-CoV-2 isolate TGR/NY/20 for 1 h at 37 °C. Following incubation of serum and virus, 50 µl of a cell suspension of Vero cells was added to each well of a 96-well plate and incubated for 48 h at 37 °C with 5% CO_2_. The cells were fixed and permeabilized as described above, and subjected to IFA using a rabbit polyclonal antibody (pAb) specific for the SARS-CoV-2 nucleoprotein (N) (produced in Dr. Diel’s laboratory) followed by incubation with a goat anti-rabbit IgG (goat anti-rabbit IgG, DyLight^®^594 Conjugate, Immunoreagent Inc.). Unbound antibodies were washed from cell cultures by rinsing the cells PBS, and virus infectivity was assessed under a fluorescence microscope. Neutralizing antibody titers were expressed as the reciprocal of the highest dilution of serum that completely inhibited SARS-CoV-2 infection/replication, after three independent VN assay performed. Fetal bovine serum (FBS) and convalescent human serum (kindly provided by Dr. Elizabeth Plocharczyk, Cayuga Medical Center [CMC]; under CMC’s Institutional Review Board protocol number 0420EP) were used as a negative and positive controls, respectively.

## Acknowledgments

The authors thank clinical veterinarian, Dr. Rebecca Cox and animal caretakers, Tiffany Williams, Kolby Stallman, Derek Vermeer and Robin Zeisneiss for excellent animal care and Patricia Federico for excellent technical assistance. Antibody reagents to SARS-CoV-2 and the virus neutralization assay used to assess serological responses were developed with support from the Cornell Feline Health Center. Mention of tradenames or commercial products is solely for the purpose of providing specific information and does not imply recommendation of endorsement by the US Department of Agriculture.

## Supplementary figures

**Supplementary Fig 1.**
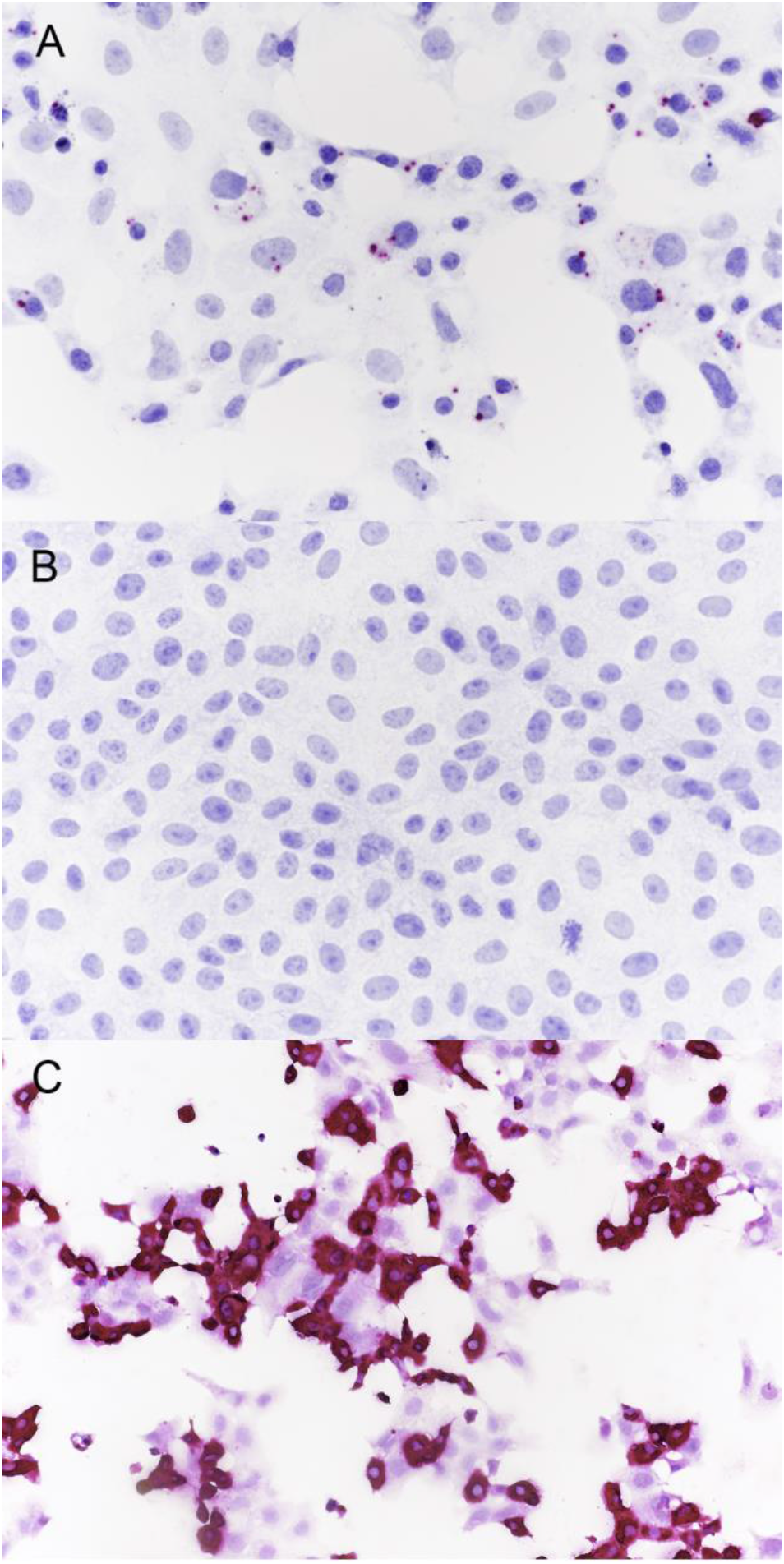
Vero cells infected with SARS-CoV-2 (A and C) or non-infected Vero cells (B) and examined 24 hrs later. Positive detection of viral anti-genomic RNA, indicative of replicating virus, is indicated by magenta staining (A). Specificity of the probe is confirmed by the lack of staining observed in the non-infected cells (B). Positive detection of accumulation of viral genomic RNA is indicated by the intense magenta staining in cell culture (C). ISH-RNAscope

**Supplementary Fig 2.**
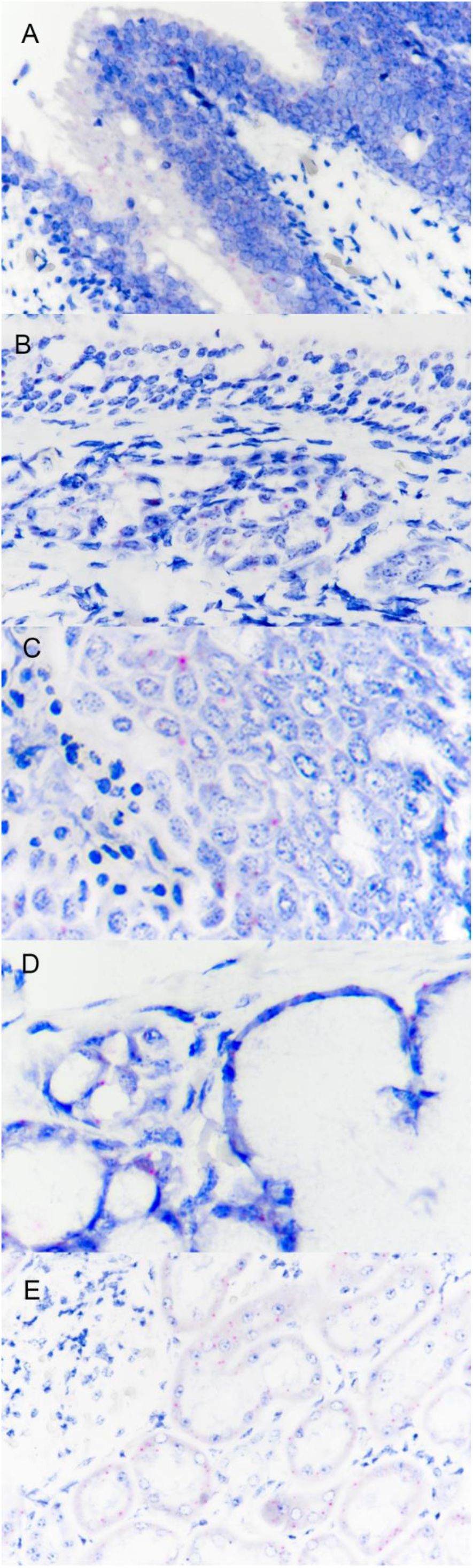
*In situ* hybridization using BaseScope technology was used to detect mRNA for the *O. virginianus* ACE2 receptor. (A and B) In nasal turbinates, positive labeling for the ACE2 receptor was detected at low levels within nasal epithelial cells and cells associated with submucosal glands. (C) Within the palatine tonsil, labeling for the receptor was seen in keratinized and non-keratinized tonsillar epithelium, including regions of lymphoepithelium overlying follicles, where the epithelial layer was heavily infiltrated by lymphoid cells. In addition, labeling was also observed in interstitial and mucous cells associated with submucosal glands (D). (E) Within the kidney, labeling was observed in proximal tubular epithelial cells, but not glomeruli, distal tubules, or collecting ducts. No ACE2 labeling was detected in lung, medial retropharyngeal lymph nodes or tracheobronchial lymph nodes. Sections of palatine tonsil (A & C), nasal turbinate (B & D) and kidney (E) from white-tailed deer fawns inoculated intranasally with SARS-CoV-2 and examined 21 days (tonsil, kidney) or 8 days (turbinate) later. Note distinct punctate magenta labeling of ACE2 receptor mRNA in tonsillar and turbinate epithelium and submucosal glands and renal tubular epithelial cells. ISH-Basescope.

**Supplementary Fig 3.**
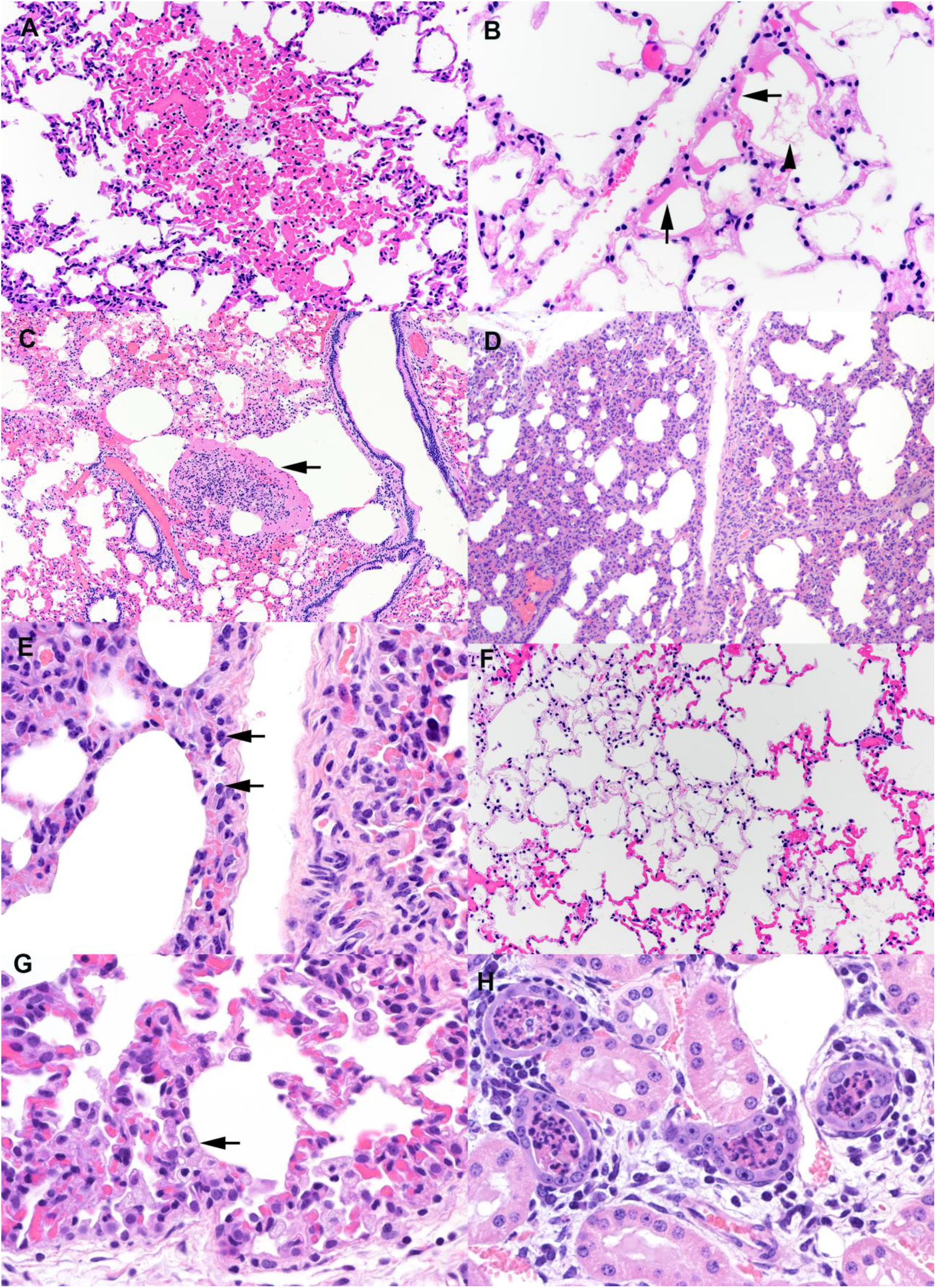
Sections of lung from white-tailed deer fawns intranasally inoculated with SARS-CoV-2 and examined 21 days later. A) Note the well demarcated focus of congestion. Alveolar septal capillaries are engorged with blood surrounded by normal appearing alveolar septa. HE B) Multiple alveolar septa are lined by bands of eosinophilic hyalinized proteinaceous material (arrows) consistent with hyaline membranes. Multiple alveoli contain flocculent to fibrillar eosinophilic material (arrow head) consistent with fibrin. HE. C) Expanded alveolus contains a large collection of fibrin, inflammatory cells, and cell debris (arrow). HE. D) Alveolar septa are expanded by an inflammatory infiltrate (interstitial pneumonia) composed primarily of lymphocytes (arrows) and macrophages (E). HE. F) Within a field of congested alveolar septa are irregular regions characterized by hypocellular septa containing few erythrocytes. Septal stroma is fibrillar and lightly eosinophilic. Multiple alveoli within these regions contain flocculent strands of fibrin. HE. G) There is type II pneumocyte hyperplasia and an increase in alveolar macrophages (arrow). HE. H) Lumens of cortical tubules in the kidney are filled with necrotic cellular debris. Renal tubules are variably lined by attenuated epithelium, occasionally have hypereosinophilic cytoplasm and pyknotic nuclei (degeneration and necrosis), and overall exhibit increased cytoplasmic basophilia (regeneration). Tubules are divided by interstitial edema and a cellular infiltrate composed of lymphocytes, plasma cells, and fewer macrophages. HE.

